# Phase-resolved transcriptomic bottlenecks in peptide cancer vaccine response

**DOI:** 10.64898/2026.08.21.746349

**Authors:** Corey Keith Goldman

**Author notes:** Corresponding author: Corey Keith Goldman, MD, PhD; MedAlliance Medical Health Services, 625 East Fordham Road, Bronx, NY 10458, United States of America.

## Abstract

**Background:** Peptide cancer vaccines can elicit antigen-specific immunity, but peripheral immunogenicity often does not translate into durable tumor control.

**Methods:** We analyzed transcriptomic profiles across three public human peptide-vaccine cohorts: C1/GSE278476, a MUC1 plus Poly-ICLC PBMC RNA-seq cohort with ordered anti-MUC1 IgG response classes; C2/GSE85698, manufactured dendritic-cell vaccine preparations linked to TARP ELISpot response; and C3/GSE53922, baseline PBMC expression linked to overall survival after personalized peptide vaccination in castration-resistant prostate cancer. Prespecified gene modules were summarized as mean standardized scores and tested with endpoint-appropriate cohort-level models with within-family FDR control.

**Results:** Baseline immune-readiness was favorable in C1 (beta=0.301, p=0.0072, q=0.093; permutation p=0.0088) and associated with longer survival in C3 (HR=0.662, 95% CI 0.532-0.823, p=0.000206, q=0.00126). Baseline erythroid/inflammatory drag showed the opposite direction in C1 (beta=-0.258, p=0.031, q=0.202; permutation p=0.0324) and was associated with inferior survival in C3 (HR=1.390, 95% CI 1.181-1.636, p=0.0000755, q=0.000982). In C2, lower tolerogenic/myeloid dendritic-cell product-state expression was observed in strong ELISpot responders (8/19 focused genes q<0.05). At C1 week 2, a priming/costimulation/mTOR-AKT module showed an FDR-significant cross-sectional association with response class (permutation p=0.0026), but paired within-person change was not significant.

**Conclusions:** Public peptide-vaccine transcriptomic data support a phase-linked model in which host readiness, erythroid/inflammatory drag, dendritic-cell product state, and early priming are measurable response-linked layers. These retrospective cohorts do not establish causality, biomarker status, clinical utility, or durable tumor control.

## Introduction

Cancer vaccines represent one of the oldest therapeutic strategies in immuno-oncology [1,2]. The conceptual premise is straightforward: present tumor-associated antigens in an immunologically favorable context, generate antigen-specific T cells, and direct those cells to eliminate tumor.

Decades of clinical experience confirm that peptide vaccines can generate measurable antigen-specific immune responses in a meaningful fraction of patients [1,2]. What they have not reliably generated is durable tumor control. The disconnect between peripheral immunogenicity and clinical benefit remains the central unsolved problem of therapeutic cancer vaccination.

The index immune-response cohort in this study used a Mucin 1 (MUC1) peptide plus Poly-ICLC vaccine. MUC1 is a well-developed tumor-associated antigen with extensive vaccine-trial experience, making GSE278476 a useful index dataset because it provides baseline and week-2 peripheral blood mononuclear cell (PBMC) transcriptomics linked to ordered immune-response classes [3–6,10,11]. The same analytic framework was then applied to a T-cell receptor gamma alternate reading frame protein (TARP) manufactured dendritic-cell vaccine preparation cohort and a personalized peptide-vaccine castration-resistant prostate cancer (CRPC) cohort. Across these cohorts, antigen identity serves as context, whereas the central question is whether phase-linked transcriptomic features can help distinguish responders from nonresponders.

Successful peptide vaccination is biologically contingent on at least four linked phases: (1) baseline host immune substrate, including a competent memory and naive T-cell compartment, functional antigen-presenting-cell machinery, and the absence of dominant immunosuppressive signals; (2) vaccine-product or antigen-presentation state, in which antigen must be delivered in a pro-immunogenic rather than tolerogenic context and, for dendritic-cell vaccines, the manufactured product must reach the injection site in an activated rather than quiescent or myeloid-biased state; (3) early post-vaccine priming and activation, during which clonal expansion and differentiation convert precursor cells into antigen-specific effectors; and (4) downstream execution and durability, in which effector cells must traffic to tumor, function within a suppressive microenvironment, and persist long enough to reduce tumor burden. Failure or attenuation at any phase can prevent clinical benefit despite measurable activity at earlier steps [1,2].

Transcriptomic profiling of peripheral blood and manufactured cell products provides a practical route to interrogate these sequential phases in clinical cohorts. Gene expression data from peripheral blood mononuclear cells (PBMCs) can capture host immune composition and activation state before and after vaccination without requiring invasive tumor sampling.

Transcriptional profiling of manufactured dendritic-cell vaccine preparations can assess product quality independently of downstream patient biology [7]. Baseline expression data linked to survival in independent cohorts can test whether host-state signals have clinical-outcome relevance beyond immunogenicity alone [8,9]. What has been missing is a coordinated analysis of these complementary evidence streams within a consistent biological framework.

This study applied a prespecified phase-resolved transcriptomic framework across three public peptide vaccine cohorts to map recurring associations aligned with biological response phases. The analysis focuses on three measurable domains with consistent evidence: (1) baseline host immune state, including immune readiness relative to erythroid/inflammatory drag; (2) dendritic-cell product tolerogenic/myeloid state; and (3) early post-vaccine PBMC priming and costimulation activity. By contrast, phase 4 - downstream execution and durability, including tumor trafficking, intratumoral function, disease response, and persistence - remains unresolved because no public human peptide-vaccine transcriptomic cohort links transcriptomic state, antigen-specific immune response, disease response, and survival within the same analytic dataset.

## Methods

### Study design and analytic objective

This study used a phase-resolved transcriptomic framework to map immune-response biology across three public peptide vaccine datasets with distinct but complementary endpoint structures. The analytic objective was to determine whether prespecified gene modules, each representing a discrete biological phase of vaccine response, showed directionally consistent associations across cohorts when tested within their respective endpoint partitions. An evidence-grading schema distinguished primary signals (false-discovery rate (FDR)-significant within prespecified analysis families), supportive signals (nominally significant or biologically coherent with primary findings), and absent signals (no detected association). Absent signals are treated as informative, identifying phases that current public data cannot resolve.

### Cohort identification and analytic roles

Candidate datasets were identified by structured searches of public repositories including the Gene Expression Omnibus (GEO), ArrayExpress/BioStudies, and trial-linked registries through May 5, 2026. Public cancer-vaccine transcriptomic resources were sparse and fragmented: adjacent datasets were available for host-state profiling, manufactured vaccine-product state, selected immune-response readouts, or clinical outcome, but no public human peptide-vaccine transcriptomic cohort identified in the search linked all four layers in the same patients. Datasets were retained as analytic cohorts when they contained transcriptomic data linked to an immune-response, product-state, or clinical-outcome endpoint with sufficient metadata for covariate-aware analysis. Three cohorts were selected, each assigned to the endpoint category supported by its available data structure (Table 1).

**Table 1.** Analytic cohorts, endpoints, and manuscript role.

| Cohort | Specimen & context | Endpoint | Sample size | Manuscript role | Endpoint scope & limits |
| --- | --- | --- | --- | --- | --- |
| C1 / GSE278476 | PBMC RNA-seq; MUC1 peptide + Poly-ICLC vaccine; baseline and week 2 | Ordered immune-response class: NR = nonresponder; LR = low responder; HR = high responder | 89 PBMC profiles; 69 donors; BL n=46; Wk2 n=43; paired BL/Wk2 subset available | Primary circulating immune-response cohort | Blood-based readout of host state and week-2 peripheral activity; clinical benefit and tumor engagement not directly measured |
| C2 / GSE85698 | Manufactured DC-vaccine product; TARP peptide vaccine product expression arrays | Strong ELISpot responder vs weak/nonresponder; continuous ELISpot secondary | 18 product samples: 4 strong responders, 14 comparators (after replicate handling) | DC-vaccine/APC state and priming-potency cohort | Product-state readout linked to ELISpot; TME and clinical outcome not measured |
| C3 / GSE53922 | Baseline PBMC expression; personalized peptide vaccination; CRPC | Overall survival | 112 baseline PBMC samples; 87 OS events; age available for 104 | Clinical-outcome corroboration cohort | Survival-associated blood signal; immune-response class and vaccine mechanism not directly measured |

Abbreviations: APC, antigen-presenting cell; BL, baseline; CRPC, castration-resistant prostate cancer; DC, dendritic cell; ELISpot, enzyme-linked immunospot; HR, high responder; LR, low
responder; NR, nonresponder; OS, overall survival; PBMC, peripheral blood mononuclear cell; TME, tumor microenvironment; Wk2, week 2.

C1 (GSE278476) comprised 89 peripheral blood mononuclear cell (PBMC) RNA-seq profiles from 69 donors enrolled in a MUC1 peptide plus Poly-ICLC adjuvant vaccine trial [10,11].

Immune-response class was defined as nonresponder (NR), low responder (LR), or high responder (HR) and encoded numerically as NR=0, LR=1, and HR=2 for ordered trend modeling. This coding preserved the biological response hierarchy while allowing module scores to be tested for monotonic trends across response classes. Paired baseline and week-2 samples were available from a subset of donors.

C2 (GSE85698) comprised manufactured dendritic-cell vaccine-preparation expression arrays from a T-cell receptor gamma alternate reading frame protein (TARP) peptide vaccine trial [7,12]. After handling of product-level and technical replicates, 18 product samples from 4 strong enzyme-linked immunospot (ELISpot) responders and 14 weak responders or nonresponders were retained. C2 evaluated manufactured antigen-presenting-cell (APC) vaccine-state biology, not donor PBMCs or post-injection activity.

C3 (GSE53922) comprised 112 baseline peripheral blood mononuclear cell (PBMC) expression profiles from patients enrolled in a personalized peptide vaccination trial in castration-resistant prostate cancer (CRPC) [8,9]. Overall survival data were available for 87 patients with events; age data were available for 104. C3 served as the independent clinical-outcome corroboration cohort.

### Endpoint-coverage framework

Peptide vaccine outcomes were organized using a hierarchical endpoint-coverage framework distinguishing confirmed durable clinical benefit (P1-P2), immune response without documented clinical improvement (P3, P6), no response (P4), and ambiguous or partially characterized patterns (P5, P7-P8). C1 and C2 populate partition P6 (transcriptomic data with immune-response endpoints; clinical benefit not linked). C3 populates P7 (survival endpoint; antigen-specific immune-response data not linked). The integrated partitions that link transcriptomic state, antigen-specific immune response, disease response, and survival remain unpopulated in current public data and are addressed as a central evidence gap in the Discussion.

### Module construction and scoring

Immune-response modules were constructed from prespecified gene sets representing eight biological phases: (1) immune readiness and memory substrate, (2) erythroid/inflammatory drag, (3) antigen presentation and innate sensing, (4) priming, costimulation, and mechanistic target of rapamycin (mTOR)-AKT signaling, (5) cytotoxic support and killing potential, (6) trafficking and tumor-entry readiness, (7) suppression and exhaustion, and (8) durability and persistence. For main-text presentation, these were grouped into four biological domains: baseline host state, DC-vaccine/APC state, early post-vaccine priming and activation, and downstream execution and durability. The overall study architecture and evidence framework are summarized in Fig 1.

**Fig 1.**
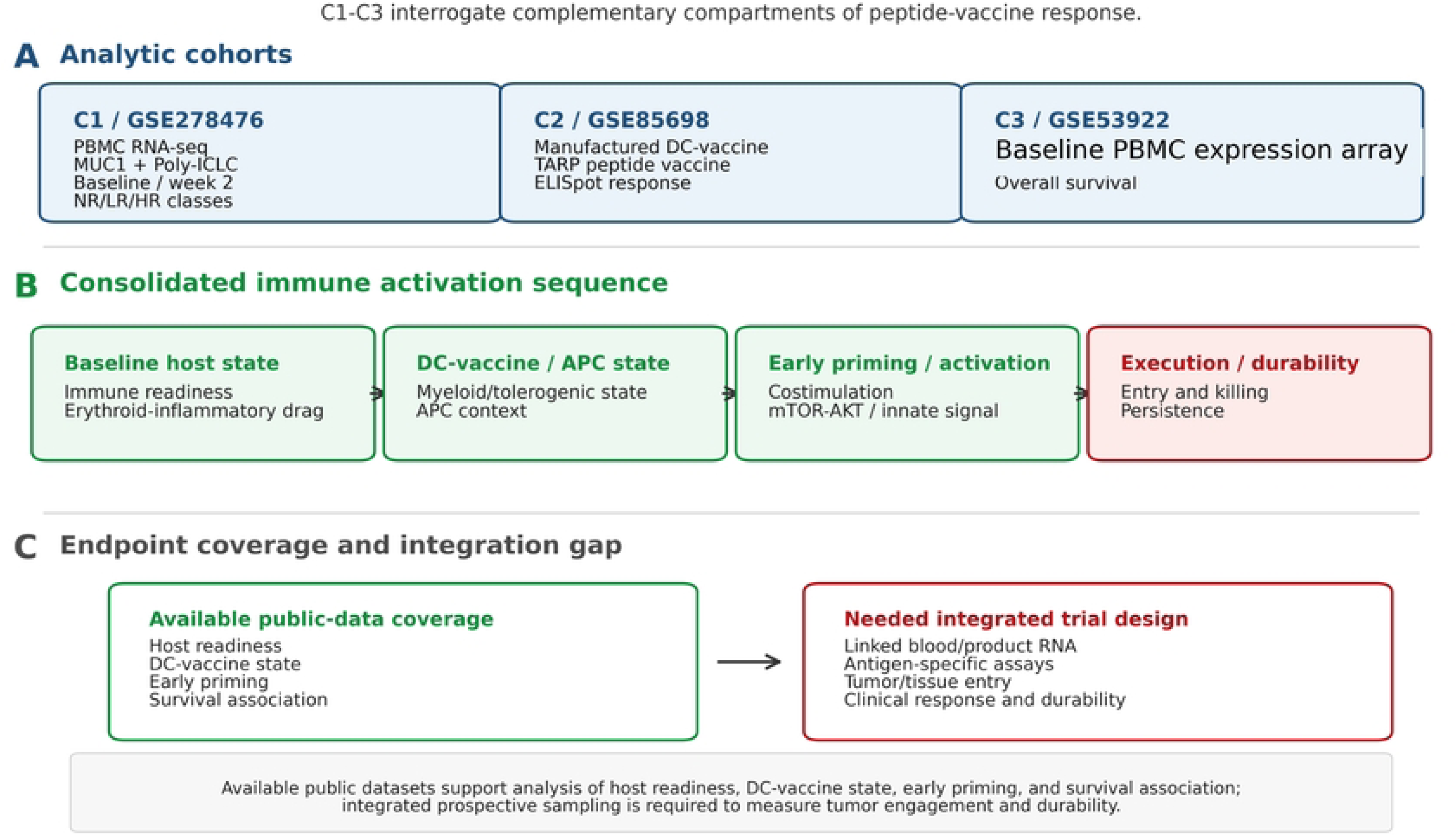
Study architecture and phase-resolved evidence framework. Three analytic cohorts were mapped to complementary measurable compartments of peptide-vaccine response. C1/GSE278476 provides the index MUC1 plus Poly-ICLC peripheral blood mononuclear cell (PBMC) RNA-seq cohort at baseline and week 2, linked to ordered immune-response classes (NR, nonresponder; LR, low responder; HR, high responder). C2/GSE85698 evaluates manufactured dendritic-cell vaccine preparations linked to TARP peptide ELISpot response and is interpreted as dendritic-cell vaccine/APC-state biology. C3/GSE53922 provides baseline PBMC expression-array associations with overall survival after personalized peptide vaccination in CRPC. Prespecified modules were consolidated into four manuscript-facing domains: baseline host state, dendritic-cell vaccine/APC state, early priming/activation, and execution/durability. Green domains and callouts indicate compartments supported by available public transcriptomic cohorts; the red downstream domain indicates the unresolved tumor-engagement and durability gap requiring integrated prospective sampling.

Primary module scoring used a mean standardized gene-module score: for each cohort/platform, measured expression values were standardized within cohort and then averaged across the available genes in each prespecified module to generate a per-sample module score. Scores were standardized to z-scores within each cohort before regression modeling. Rank-based enrichment scores were retained as sensitivity or proxy analyses where explicitly labeled, particularly for erythroid-lineage checks. HPCA- and Blueprint-reference analyses were used as orthogonal enrichment/proxy checks rather than absolute cell-fraction estimates [14,15]. Complete module gene sets, internal identifiers, and overlap matrices are provided in Supplementary Tables S1 and S5.

### Statistical analysis

C1 module scores were tested against ordered immune-response class using covariate-adjusted linear-trend models, with response encoded as NR=0, LR=1, and HR=2 (NR, nonresponder; LR, low responder; HR, high responder). Sex and batch were prespecified covariates. Baseline analyses used 46 samples (HR=9, LR=4, NR=33); week-2 analyses used 43 samples (HR=19, LR=5, NR=19). A paired-sample delta analysis (week 2 minus baseline module score) was performed in donors with both timepoints to distinguish within-person induction from response-associated cross-sectional state. Covariate-adjusted Freedman-Lane permutation sensitivity tests were performed for the main C1 signals.

C2 analyses used exact two-sided Mann-Whitney/rank tests and Cliff’s delta comparing strong ELISpot responders (n=4) to weak responders/nonresponders (n=14) across the prespecified 19-gene focused tolerogenic/myeloid DC-vaccine state panel, with Benjamini-Hochberg FDR control across the focused panel. Continuous ELISpot sensitivity analyses were performed as a secondary check. C3 analyses used Cox proportional hazards regression of standardized module scores against overall survival, with age-adjusted models as the prespecified primary sensitivity analysis. Custom Schoenfeld residual trend diagnostics were generated as supplementary audit checks and were not used as confirmatory tests.

Benjamini-Hochberg false-discovery rate (FDR) correction was applied within prespecified cohort-domain analysis families rather than globally across all cohorts [16]. This cohort-level approach was chosen because C1, C2, and C3 addressed non-exchangeable biological questions using different specimen types, endpoints, and statistical models: ordered immune-response class in PBMCs, ELISpot-linked manufactured dendritic-cell vaccine preparation state, and overall survival from baseline PBMCs. A single global FDR correction across these heterogeneous analyses would pool tests that do not represent one statistical family and would obscure the prespecified endpoint-specific inference. Within-family FDR therefore controlled multiplicity within each interpretable testing family while preserving the biological meaning of each cohort’s supported endpoint partition. Relative-path manuscript-level reproducibility scripts, model diagnostics, processed supporting tables, and FDR family definitions are provided in the supplementary methods audit files.

### Overlap-aware module consolidation and redundancy sensitivity analysis

Because several modules shared canonical immune genes, overlap-aware module consolidation was performed as a sensitivity analysis to evaluate whether principal findings reflected shared gene membership rather than separable biological programs. Shared genes were assigned to the module with the strongest prespecified biological rationale, and residual gene sets were rescored as nonredundant module scores. Primary results are reported from the original prespecified modules, with nonredundant sensitivity scores used to determine whether a signal remained independently interpretable. Signals that materially attenuated after consolidation were classified as not independently resolved and mapped to the downstream execution/durability gap pending tissue-level or longitudinal confirmation.

Manuscript-level reproducibility checks were implemented in relative-path Python 3 scripts using deterministic calculations and are supplied with the supporting tables. Public raw/source data remain available from GEO; processed module-score tables and focused patient-level C2 expression values are included as Supporting Information.

### Ethics statement

This study is a secondary analysis of de-identified, publicly available datasets and did not involve new interaction with human participants or access to identifiable private information by the author. No additional institutional review board approval or participant consent was required for this secondary analysis.

### Artificial intelligence tools and technologies

AI-assisted tools, including OpenAI ChatGPT/Codex, were used to support literature organization, code drafting, figure-format checking, manuscript editing, and PLOS ONE reformatting. The author reviewed, verified, and approved all analyses, interpretations, citations, figures, tables, and final manuscript text, and remains responsible for the accuracy and integrity of the work.

## Results

C1, C2, and C3 were selected because they span complementary measurable compartments of peptide vaccine response: circulating immune state before and shortly after vaccination (C1), manufactured APC/DC-vaccine quality (C2), and baseline blood state linked to long-term clinical outcome (C3). Their heterogeneity in antigen, platform, tumor setting, and endpoint is a feature of the design rather than an attempt at direct quantitative pooling; the analysis asks whether phase-aligned signals recur across disparate vaccine contexts. Current public data most strongly populate immune-response-linked and product-state compartments (C1 and C2) and a clinical-outcome-linked compartment (C3), whereas integrated compartments linking transcriptomic state, confirmed antigen-specific immune response, disease response, and survival remain unpopulated.

### Baseline blood immune-readiness and erythroid/inflammatory drag show opposing associations with immune-response class in C1

C1 baseline results are displayed in Fig 2. Among the 46 baseline PBMC profiles (HR=9, LR=4, NR=33), higher immune-readiness module score was positively associated with stronger ordered immune-response class, defined for C1 as NR=0, LR=1, and HR=2 (beta=0.301, p=0.0072, q=0.093). A covariate-adjusted Freedman-Lane permutation sensitivity preserved nominal support for this association (permutation p=0.0088). The readiness module is anchored by genes marking naive and central memory T cells (IL7R, TCF7, LEF1, CCR7, SELL) and lymphoid survival and priming competence (BCL2, LTB, FOXO1), reflecting a blood immune substrate positioned for antigen-driven expansion [17–19].

**Fig 2.**
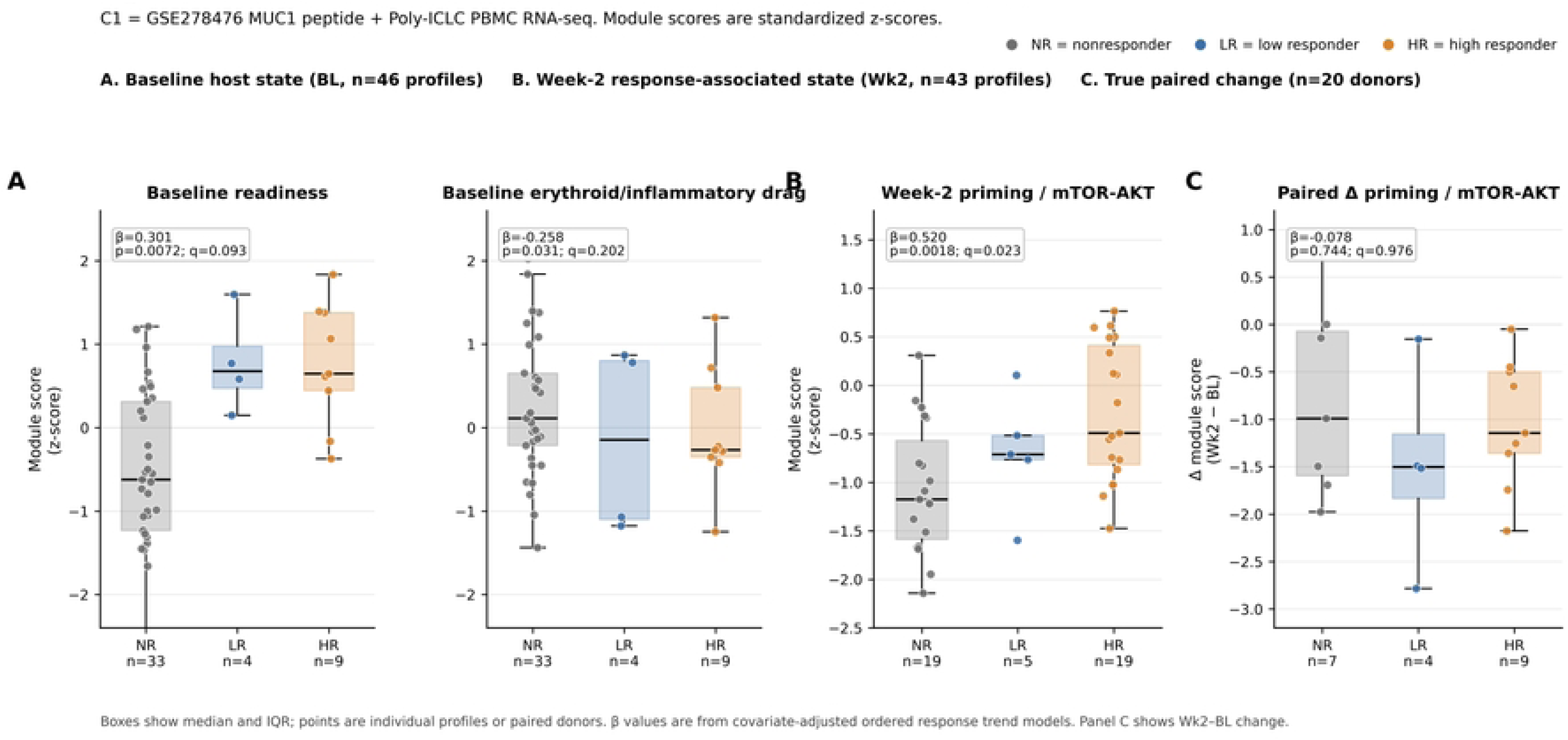
C1 PBMC immune-response architecture across response classes. (A) Baseline module z-scores stratified by ordered immune-response class (NR, nonresponder; LR, low responder; HR, high responder; C1 trend encoding NR=0, LR=1, HR=2; n=46 profiles). Immune-readiness score increases directionally with response class, whereas erythroid/inflammatory-drag score shows the reciprocal direction. These baseline C1 associations are supportive and FDR-attenuated. (B) Week-2 priming/costimulation/mTOR-AKT module score stratified by response class (n=43 profiles), showing an FDR-significant cross-sectional association. (C) True paired within-person change in the same priming module among donors with both baseline and week-2 samples (n=20 donors), showing no significant paired-delta association. Boxes show median and interquartile range; points indicate individual profiles or paired donors. Beta values are from covariate-adjusted ordered-response trend models.

The erythroid/inflammatory-drag module showed the reciprocal pattern: higher baseline drag score was associated with lower immune-response class (beta=-0.258, p=0.031, q=0.202), with nominal support retained in covariate-adjusted permutation sensitivity testing (permutation p=0.0324). The erythroid component of this signal -- driven by ALAS2, AHSP, GYPA, HBB, SLC4A1, KLF1, and GATA1 -- is consistent with stress erythropoiesis and myeloid-erythroid skewing in circulating blood [20–22]. These processes are biologically aligned with tumor-associated stress hematopoiesis, which can expand erythroid-myeloid programs and suppressive immature myeloid populations while impairing dendritic-cell differentiation [20–24]. The S100A8/S100A9 component (calprotectin) adds a chronic inflammatory and well-described myeloid-derived suppressor cell (MDSC)-associated alarmin tone [23,24]. Erythroid-lineage proxy/enrichment checks using HPCA and Blueprint references supported this interpretation, arguing against a signal driven solely by PBMC isolation artifact [14,15].

Because these baseline PBMC RNA profiles were obtained before vaccine administration and before response classification, the C1 host-state modules are temporally positioned as candidate predictors of subsequent anti-MUC1 immune-response class. This use of predictor is limited to temporal and statistical prediction; the present analysis does not establish causality, treatment-effect modification, a validated biomarker, or clinical decision utility.

### Week-2 priming and costimulation is the leading post-vaccine PBMC signal, but within-person induction is not confirmed

Week-2 C1 results are shown in Fig 2. The priming/costimulation/mTOR-AKT module score at week 2 was the strongest cross-sectional association in the full C1 analysis (n=43; HR=19, LR=5, NR=19; beta=0.520, p=0.0018, q=0.023). A covariate-adjusted permutation sensitivity preserved support for this week-2 association (permutation p=0.0026). Genes driving this signal include CD28, ICOS, CD40LG, CD40, CD27, NFKB1, RELA, MTOR, AKT1, and RPS6KB1 -- a signature of early T cell costimulation and downstream anabolic and proliferative signaling that collectively marks an activated, expanding lymphocyte state [19,25,26].

The paired-sample delta analysis, however, was null: in donors with both baseline and week-2 samples, the week-2 minus baseline priming/costimulation delta showed no significant association with immune-response class (β=-0.079, p=0.744). This means that the week-2 cross-sectional result reflects a difference in transcriptional state between high responders and comparators at week 2, but does not prove that the vaccine induced this state within individual donors. High responders arrive at week 2 in a more activated priming state; whether this reflects vaccine-driven induction, pre-existing biological differences that become more apparent post-vaccination, or both cannot be resolved without a properly powered within-person design. Thus, the week-2 priming score should be interpreted as an early response-state classifier, not as mechanistic proof of within-person vaccine induction. This distinction is critical for monitoring protocol design: prospective trials should include prespecified within-person delta analysis with adequate sample size to separate induction from response-associated state.

### Tolerogenic and myeloid-lineage gene expression in the manufactured DC-vaccine state is associated with ELISpot response failure in C2

C2 focused DC-vaccine state results are shown in Fig 3. Strong ELISpot responders had lower expression of tolerogenic and myeloid-lineage genes in their manufactured DC-vaccine preparations relative to weak responders and nonresponders. Using exact patient-level rank testing with Benjamini-Hochberg correction across the 19-gene focused panel, eight genes reached FDR q<0.05, all lower in strong responders: IL10, CD163, MMP14, TGFBI, S100A12, CD14, S100A9, and S100A8 (Fig 3). TGFBI (TGF-beta-induced) is distinct from TGFB1, which was included in the focused panel but was not FDR-significant. The pattern is consistent with reduced tolerogenic/myeloid polarization rather than mature, immunogenic dendritic-cell differentiation. Binary ELISpot classification sensitivity analysis supported the same broad lower-expression direction.

**Fig 3.**
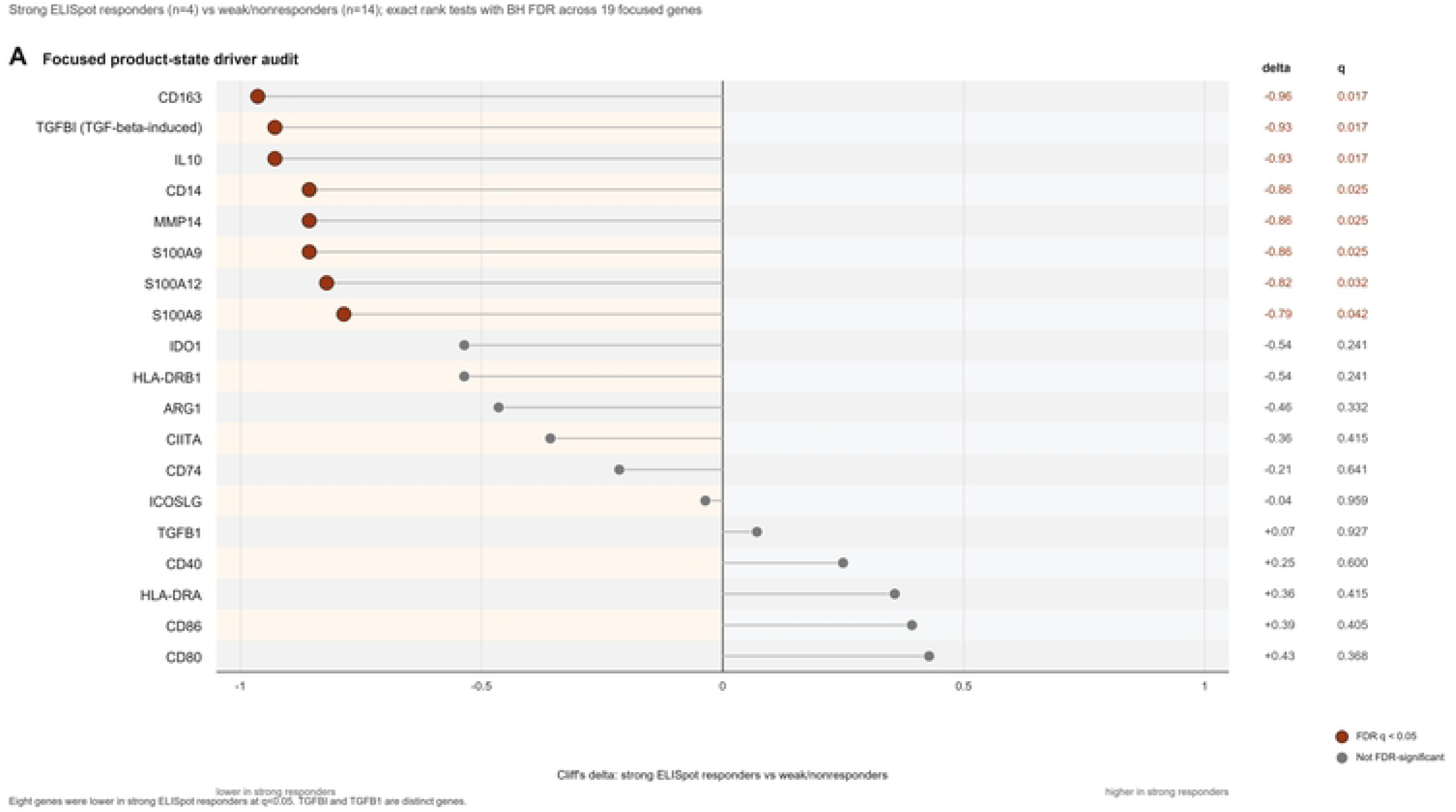
C2 manufactured dendritic-cell vaccine-state focused tolerogenic/myeloid panel. Ranked dot/effect-size plot of the 19-gene focused tolerogenic/myeloid dendritic-cell vaccine-state panel comparing strong ELISpot responders (n=4) with weak responders and nonresponders (n=14). Effect size is plotted as strong-responder minus comparator expression; negative values indicate lower product-gene expression in strong responders. Dark red points indicate focused-panel genes significant at exact rank-test Benjamini-Hochberg FDR q<0.05; gray points indicate non-significant genes. Lower expression of IL10, CD163, MMP14, TGFBI, S100A12, CD14, S100A9, and S100A8 supports a lower tolerogenic/myeloid DC-vaccine state in strong ELISpot responders. TGFBI and TGFB1 are distinct genes. Because the strong-responder group is small, this result is interpreted as hypothesis-generating product-state biology rather than a validated product-release biomarker.

Because C2 profiles the manufactured product rather than donor blood, the result identifies a candidate product-state readout linked to ELISpot priming potency [7]. The C2 cohort is small (n=4 strong responders, n=14 comparators). Leave-one-strong-responder sensitivity showed that the exact FDR-significant gene list is not stable to omission of individual strong responders, although the broad lower tolerogenic/myeloid direction was largely preserved. These results are therefore hypothesis-generating and should not be interpreted as a qualified product-release assay. They nonetheless support prospective testing of tolerogenic gene expression profiling as a candidate product-quality measure, complementary to standard phenotypic release markers.

### Baseline immune-readiness and erythroid/inflammatory drag are associated with overall survival in C3

C3 survival results are shown in Fig 4. In 112 baseline PBMC profiles from a personalized peptide vaccine trial in CRPC - an independent cohort in a different tumor type with a different endpoint structure - the baseline immune-readiness module score was significantly associated with longer overall survival (HR=0.662 per 1 SD increase, 95% confidence interval (CI) 0.532-0.823, p=0.000206, q=0.00126). The association persisted after age adjustment (HR=0.675, q=0.00815).

**Fig 4.**
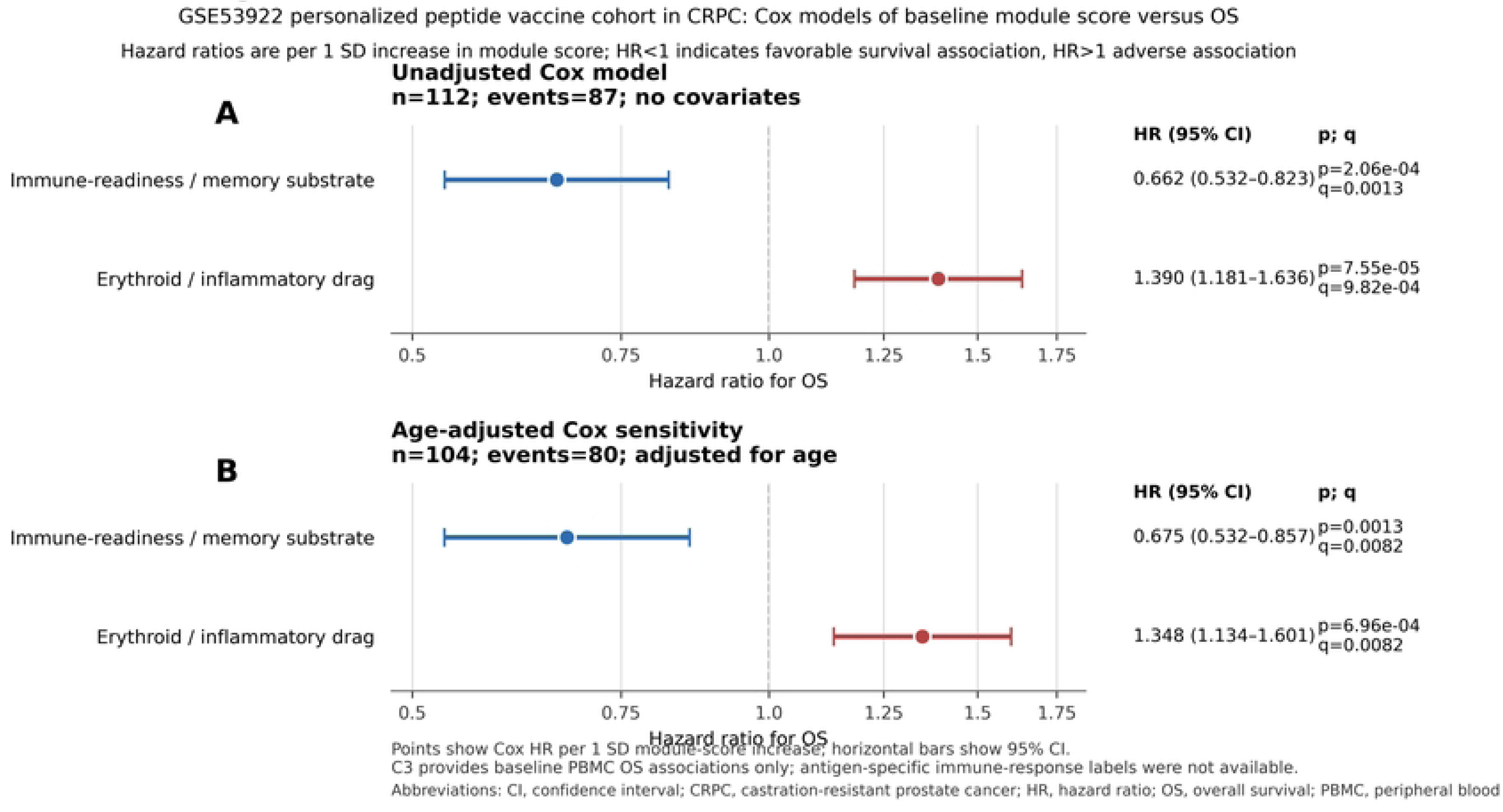
C3 baseline PBMC module score associations with overall survival. Forest plots show Cox proportional-hazards models for prespecified baseline PBMC host-state modules in GSE53922, a personalized peptide vaccine cohort in CRPC. (A) Unadjusted models. (B) Age-adjusted sensitivity models. Hazard ratios are shown per 1-SD increase in module score; horizontal bars indicate 95% confidence intervals. Higher immune-readiness/memory-substrate score was associated with lower mortality risk, whereas higher erythroid/inflammatory-drag score was associated with higher mortality risk. Age-adjusted models are shown as sensitivity analyses because age was available for this cohort. Abbreviations: CI, confidence interval; CRPC, castration-resistant prostate cancer; FDR, false-discovery rate; HR, hazard ratio; OS, overall survival; PBMC, peripheral blood mononuclear cell.

Higher erythroid/inflammatory-drag score was significantly associated with shorter overall survival (HR=1.390 per 1 SD increase, 95% CI 1.181-1.636, p=0.0000755, q=0.000982; age-adjusted HR=1.348, q=0.00815) (Fig 4). These are the most statistically robust associations in the entire analysis. The convergence of readiness-favorable and drag-adverse directionality across immune-response class in C1 and overall survival in C3 -- in distinct tumor types, vaccine platforms, and patient populations -- provides the cross-cohort corroboration central to this study’s evidence architecture. The C3 survival associations suggest that the erythroid/inflammatory drag score captures an adverse systemic host-state burden relevant to outcome, but do not establish vaccine-specific treatment-effect prediction.

Main graded evidence findings across C1-C3 are summarized in Table 2.

**Table 2.** Main graded evidence findings across C1-C3.

| Evidence domain | Cohort | Endpoint | Main statistic | Interpretation |
| --- | --- | --- | --- | --- |
| Baseline immune-readiness / memory substrate | C1 | Baseline PBMC HR/LR/NR | $\beta=0.301$; $p=0.0072$; $q=0.093$; $n=46$ | Supportive: higher readiness score associates with stronger immune-response class. Directionally consistent in the original module and nonredundant sensitivity score. Not treated as proof of pre-existing antigen-specific memory. |
| Baseline erythroid / inflammatory drag | C1 | Baseline PBMC HR/LR/NR | $\beta=-0.258$; $p=0.031$; $q=0.202$; $n=46$; ssGSEA enrichment corroborative | Supportive: higher drag associates with poorer immune-response class. HPCA/Blueprint proxy checks support erythroid-lineage direction. |
| Early priming / costimulation / mTOR-AKT | C1 | Week-2 PBMC HR/LR/NR | $\beta=0.520$; $p=0.0018$; $q=0.023$; $n=43$; paired delta $\beta=-0.079$, $p=0.744$ | Primary: strongest post-vaccine PBMC signal. Interpreted as a week-2 response-associated transcriptional state; within-person vaccine-driven induction is not confirmed by paired delta analysis. |
| Tolerogenic / myeloid DC-vaccine state | C2 | Strong ELISpot vs weak/NR | 8/19 focused panel genes lower at $q < 0.05$ in strong responders; n=4 strong, n=14 comparators | Primary C2 result: lower tolerogenic/myeloid product-state gene expression associated with stronger antigen-specific priming. Small n; treated as hypothesis-generating for prospective product-quality studies. |
| Survival-associated immune-readiness | C3 | Baseline PBMC OS | HR=0.662; 95% CI 0.532-0.823; p=0.000206; q=0.00126; age-adjusted HR=0.675; q=0.00815 | Primary: higher immune-readiness associates with favorable OS in independent peptide-vaccine outcome cohort. Age-adjusted result robust. |
| Survival-associated erythroid / inflammatory drag | C3 | Baseline PBMC OS | HR=1.390; 95% CI 1.181-1.636; p=0.0000755; q=0.000982; age-adjusted HR=1.348; q=0.00815 | Primary: higher drag score associates with adverse OS. Among the most statistically robust findings in the analysis. Replicates C1 drag directionality using an independent endpoint and tumor type. |

Abbreviations: CI, confidence interval; DC, dendritic cell; ELISpot, enzyme-linked immunospot; FDR, false-discovery rate; HR (in statistics column), hazard ratio; HR/LR/NR, high/low/nonresponder; mTOR-AKT, mechanistic target of rapamycin-protein kinase B; NR, nonresponder; OS, overall survival; PBMC, peripheral blood mononuclear cell; q, Benjamini-Hochberg false discovery rate. Evidence grade: Primary = FDR-significant within prespecified analysis family; Supportive = nominally significant or coherent with primary signals.

### Overlap-aware consolidation supports the evidence architecture

Overlap-aware redundancy results are summarized in Fig 5 and Table 3. After assigning shared canonical immune genes to their primary biological domain and recomputing nonredundant module scores, the main findings in the baseline host-state domain (readiness and drag, in both C1 and C3) and the DC-vaccine/APC-state domain (tolerogenic DC tone, in C2) retained their effect direction and nominal significance. The week-2 priming/costimulation signal in C1 also retained its leading statistical rank in the nonredundant sensitivity analysis.

**Fig 5.**
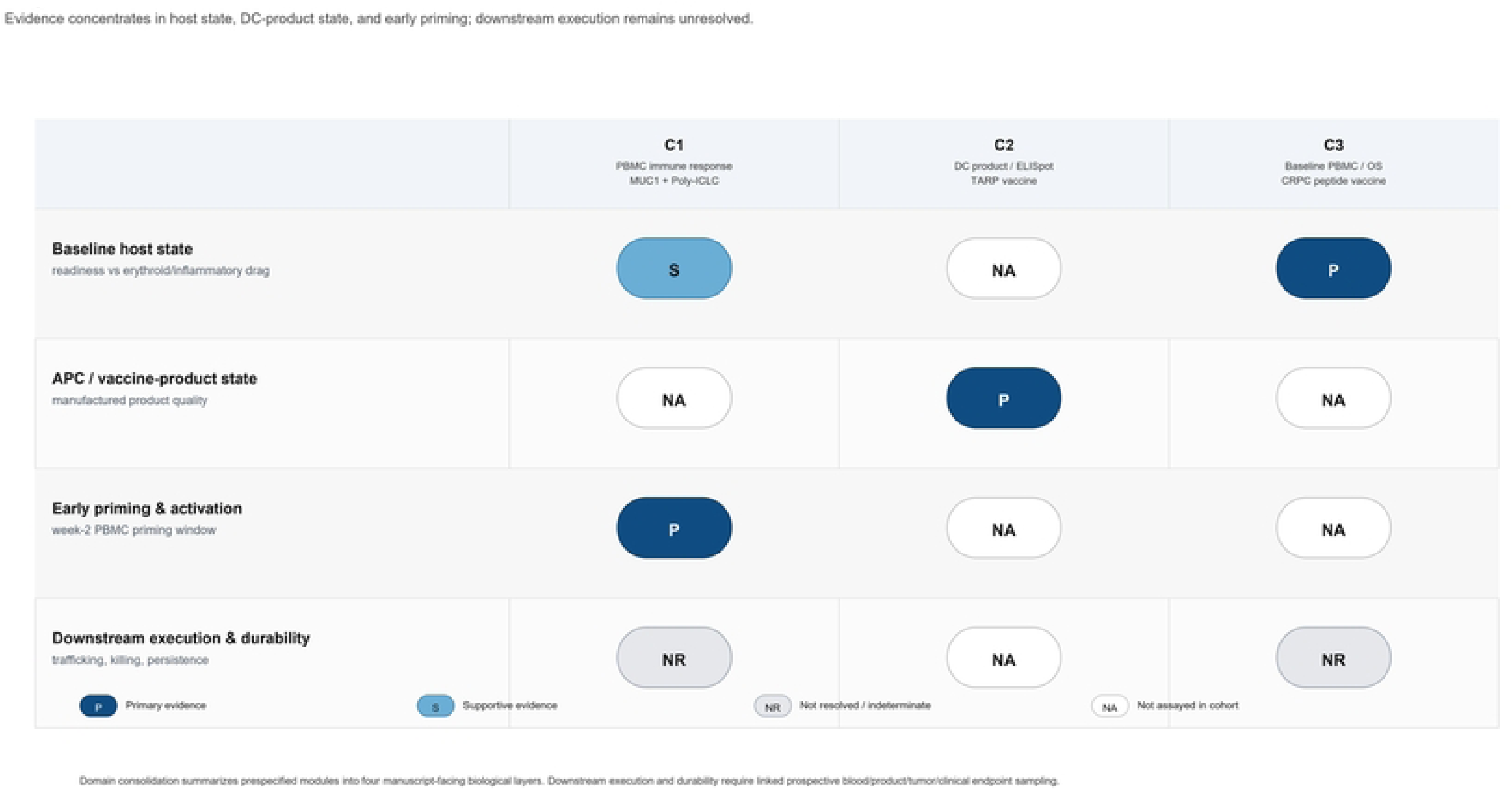
Consolidated evidence architecture across peptide-vaccine transcriptomic cohorts. Evidence-symbol map summarizing four manuscript-facing biological domains across the three analytic cohorts after domain consolidation. Dark blue indicates primary evidence, light blue indicates supportive evidence, gray indicates not resolved or indeterminate evidence, and white indicates not assayed in that cohort. P, primary evidence; S, supportive evidence; NR, not resolved/indeterminate; NA, not assayed. Evidence states were assigned according to cohort-supported endpoint availability and prespecified statistical interpretation. The map shows that available public transcriptomic data concentrate on host state, dendritic-cell vaccine state, and early priming, while downstream execution and durability remain insufficiently resolved without linked prospective blood/product/tumor/clinical endpoint sampling.

**Table 3.** Consolidated domain evidence map after module-redundancy adjudication.

| Main-text domain | Evidence source | Primary interpretation | Sensitivity / scope note |
| --- | --- | --- | --- |
| Baseline host state | C1 (immune-response class) and C3 (overall survival) | Most consistent domain. Immune-readiness is favorable; erythroid/inflammatory drag is adverse. Both associations replicate directionally across independent cohorts and distinct endpoints. | Overlap-aware redundancy results support treating durability as part of the readiness substrate unless longitudinal TCR persistence data are available. |
| APC / vaccine-product state | C2 DC-vaccine state focused panel (primary); broad suppression/tolerogenic module (corroborating context) | Lower tolerogenic/myeloid product-state gene expression associates with stronger ELISpot response. Biologically coherent with manufacturing-driven tolerogenic drift. | Focused 19-gene panel leads interpretation. Broad suppression module is corroborating, not an independent discovery. Small C2 n requires prospective replication. |
| Early post-vaccine priming and activation | C1 week-2 PBMC data | Priming/costimulation/mTOR-AKT is the leading post-vaccine PBMC response-associated program at week 2. Strongest single association in the C1 analysis. | Paired-delta analysis null: cross-sectional result reflects response-associated state, not confirmed within-person vaccine-induced induction. Prospective delta analysis required. |
| Downstream execution and durability | Supplementary module framework; future trial design | Current public blood and product data define the gap rather than resolve it. Cytotoxic support, trafficking, suppression, and durability modules are biologically plausible but not independently resolved by the available bulk blood/product transcriptomic data. | Tissue sampling, longitudinal TCR monitoring, ctDNA/imaging endpoints, and clinical-response-linked data are needed to quantify each downstream phase independently. |

Abbreviations: APC, antigen-presenting cell; DC, dendritic cell; ELISpot, enzyme-linked immunospot; mTOR-AKT, mechanistic target of rapamycin-protein kinase B; n, sample size; PBMC, peripheral blood mononuclear cell; TCR, T cell receptor.

By contrast, the innate/IFN-tone, cytotoxic-support, trafficking, suppression/exhaustion, and durability modules showed marked attenuation or loss of independent signal after overlap-aware consolidation, consistent with substantial gene overlap with the priming and readiness modules. These domains are not eliminated from the biological interpretation - they represent real and important phases of vaccine response biology - but the current public data do not support their independent quantification from bulk blood or product transcriptomics alone. In the available cohorts, C1 and C2 provide upstream immunogenicity endpoints, whereas C3 provides survival-linked outcome data. Downstream execution and durability therefore remain classified within the downstream execution/durability gap pending tissue-based profiling, longitudinal monitoring, and endpoint-linked clinical-outcome data (Table 3).

## Discussion

### Principal findings and their sequential organization

This phase-resolved analysis identifies three transcriptomic layers that recur across independent public peptide vaccine datasets with distinct endpoint structures: (1) baseline peripheral-blood host state, specifically the balance between immune readiness and erythroid/inflammatory drag; (2) dendritic-cell vaccine-product tolerogenicity; and (3) early post-vaccine priming/costimulation state. The cohorts differ intentionally in antigen, vaccine platform, tumor type, specimen, and endpoint. Directionally coherent signals across this heterogeneity are consistent with the central interpretation: peptide-vaccine performance is constrained by phase-aligned host, product, and priming states that can be measured prospectively but are not yet captured within any single public cohort.

Taken together, the C1 and C3 findings are consistent with a clinically intuitive model in which peptide vaccination is more likely to generate a productive immunologic response when administered into an immunologically favorable host state: preserved memory/lymphoid readiness, low erythroid-inflammatory drag, and an antigen-presenting platform that avoids tolerogenic myeloid polarization. In this framework, MUC1, TARP, and personalized peptide targets function as platform-specific antigens used to interrogate shared response architecture rather than as evidence that any one antigen uniquely governs the observed biology. This interpretation does not by itself justify a specific intervention, but it does provide a biologically grounded rationale for testing pre-immunization optimization strategies prospectively [1,2,20,22].

The present findings are broadly consistent with the primary cohort reports. In C1, they support prior observations linking responder status to a more favorable baseline immune substrate and stronger early activation-related transcriptional programs; in C2, they align with the association between lower tolerogenic/myeloid dendritic-cell product-state features and stronger antigen-specific response; and in C3, they are concordant with personalized peptide-vaccination literature showing that pretreatment blood biology influences outcome. The added contribution of the present study is the placement of these observations within a common phase-resolved framework, allowing shared response domains and the unresolved downstream execution gap to be interpreted across datasets.

The host-state interpretation is supported, although not proven, by prior interleukin-7 (IL-7) vaccine-adjuvant studies [28]. Pellegrini et al. reported that short-course IL-7 after vaccine induction improved antitumor responses, survival, and tumor T-cell infiltration in preclinical vaccine models [29]. Colombetti et al. showed that recombinant IL-7 after lentivector immunization increased BCL2 and expanded tumor-antigen-specific and memory CD8+ T-cell populations [30]. Early clinical studies using IL-7 gene-modified autologous tumor-cell vaccines similarly demonstrated feasibility and increased peripheral-blood tumor-reactive or cytolytic responses, although objective clinical responses were limited and inconsistent [31]. Together, these studies frame IL-7 less as a stand-alone antitumor solution than as a means of amplifying a receptive immune substrate.

The most relevant modern human precedent is the randomized phase II CYT107-after-sipuleucel-T trial in metastatic CRPC [32]. Pachynski et al. reported that recombinant human IL-7 after sipuleucel-T was well tolerated and expanded CD4+ T cells, CD8+ T cells, and CD56bright NK cells, whereas IFN-γ ELISpot responses were not significantly different from observation; antigen-specific proliferative and humoral responses improved over time in the IL-7 arm [32]. This pattern is directly relevant to our findings because the favorable readiness module includes IL7R together with memory- and homeostasis-associated genes such as TCF7, LEF1, BCL2, CCR7, SELL, NT5E, and PDPR [17–19,32]. The literature therefore supports IL-7 as an immune-substrate amplifier compatible with the readiness phenotype identified here, while remaining consistent with the larger point of this manuscript: a permissive baseline host state may improve immunogenicity, but it does not by itself guarantee downstream tumor control.

Any IL-7-directed pre-immunization strategy should therefore be treated as a prospective hypothesis rather than a present clinical recommendation.

Baseline readiness suggests one class of supportive intervention; the week-2 priming signal suggests another.

The week-2 activation state also has a plausible cytokine-support context. Its gene content spans PI3K-AKT-mTOR nodes (AKT1, MTOR, RPS6KB1), NF-κB-linked activation regulators (NFKB1, NFKBIA, TNFAIP3), costimulatory nodes (ICOS, CD40LG, CD27, CD70), and effector-memory/cytotoxic-survival markers (BCL2, PIM1, PIM2, NKG7, GZMB, PRF1), making it biologically compatible with interleukin-15 (IL-15)/IL-15 receptor-responsive immune architecture [25–27]. This is relevant because N-803 and related IL-15/IL-15 receptor agonist strategies are already being tested clinically as immune-support partners for MUC1-containing multivalent tumor-antigen vaccine platforms [33–35]. Within the framework of this manuscript, such agents are best viewed not as generic add-ons but as candidate consolidation partners to be timed after evidence of early priming. Conversely, downstream helper and durability strategies, including IL-21-axis support, would require separate readouts of Tfh, B-cell, memory, and persistence biology rather than inference from the week-2 priming score alone.

### The erythroid/inflammatory drag signal: an adverse systemic host-state burden

Among the three measurable layers, the erythroid/inflammatory drag finding deserves particular emphasis because it points to an adverse systemic host-state burden that may constrain vaccine responsiveness. In contrast to conventional vaccine biomarkers such as checkpoint receptor expression, regulatory T-cell frequency, or tumor-infiltrating lymphocyte density, the genes anchoring this module are not classic T-cell exhaustion markers or direct tumor-microenvironment readouts. Instead, they point to erythroid progenitor activity and myeloid alarmin signaling in circulating blood, suggesting that a host-level hematopoietic and inflammatory state may shape vaccine responsiveness before downstream tumor biology is even engaged.

Within this cross-cohort, phase-resolved framework, the erythroid/inflammatory-drag signal is a response-linked host-state finding requiring prospective validation.

Elevated erythroid-lineage gene expression in PBMCs (ALAS2, AHSP, GYPA, HBB, SLC4A1, KLF1, GATA1) is mechanistically interpretable as a readout of stress erythropoiesis driven by inflammatory and tumor-derived hematopoietic signals, including erythropoietin (EPO)-axis and myeloid-skewing cytokine programs. These processes can favor erythroid-myeloid output and suppressive immature myeloid populations, with reported impairment of dendritic-cell differentiation [20–24,36,37]. This competition at the level of common myeloid-erythroid progenitors is an increasingly recognized mechanism of tumor-induced immune suppression that is distinct from checkpoint-mediated T-cell exhaustion and has been reported to limit anti-PD-1/PD-L1 efficacy in mechanistic tumor models [22]. The co-elevation of S100A8 and S100A9 (calprotectin, a well-described MDSC-associated alarmin signal) extends this biology into the MDSC compartment and is consistent with the known role of immature myeloid cells as downstream effectors of stress erythropoiesis-driven immunosuppression [21–24].

Several related clinical and technical explanations should be considered together. Erythroid-lineage transcripts in PBMC RNA-seq can reflect reticulocyte or red-cell carryover, anemia-associated circulating erythroid precursors, inflammation, tumor burden, marrow stress, cachexia, prior therapy, or technical variation in cell preparation [20,21,36]. From a clinical perspective, these possibilities represent overlapping adverse host-state biology rather than mutually exclusive alternatives in the present public datasets. In this analysis, the drag association was not eliminated by sex/batch adjustment or HPCA/Blueprint-informed erythroid enrichment analysis, and the same domain associated with adverse overall survival in C3. These features reduce the likelihood that the signal is solely a processing artifact, but they do not fully distinguish biology from residual pre-analytic variation. Available C1-C3 metadata did not provide a harmonized hemoglobin level, reticulocyte count, iron-study panel, inflammatory markers, treatment-history variables, tumor-burden measures, or sample-processing variables sufficient to partition these contributors individually. Future validation should therefore pair bulk RNA with complete blood count indices, reticulocyte measures, iron/hepcidin/inflammatory biomarkers, treatment and disease-burden metadata, and cellular-resolution profiling using single-cell RNA-seq, CITE-seq, or sorted-population transcriptomics.

The C3 survival association for erythroid/inflammatory drag (HR=1.390 per 1 SD, p=7.55e-5) is one of the strongest statistical relationships in the entire dataset. This is not an immunogenicity correlation; it is a survival signal in an independent cohort measured at baseline, before any vaccination-induced immune response could be assessed. For trial design, this supports evaluating erythroid/inflammatory drag score as a candidate patient-stratification variable at enrollment, with a scientific rationale for testing whether reducing an adverse host-state burden -- through correction of anemia or iron-restricted erythropoiesis, MDSC-targeting approaches, TGF-beta pathway inhibition, or other myeloid-rebalancing strategies -- improves the effective immune substrate available for vaccination. Such interventions remain hypotheses for prospective testing, but the current data make the host-state question concrete and measurable.

### From peripheral immunogenicity to clinical benefit: the critical unmapped gap

If the first half of this Discussion identifies what current public datasets can resolve, the second half must address what they cannot. The most consequential structural finding of this analysis is an absence rather than a specific gene association: no public human peptide-vaccine transcriptomic cohort identified in our repository search currently links transcriptomic profiling to antigen-specific immune assays, biochemical or radiographic disease response, and survival within the same analytic dataset. In the available public cohorts, C1 and C2 provide upstream immunogenicity endpoints without linked survival outcomes, whereas C3 provides survival-linked outcome data without paired antigen-specific immune-response measures. As a result, downstream execution and durability cannot be mapped across the full pathway from immune priming to clinical outcome. This is not simply a limitation of the present study; it exposes a recurring monitoring problem in vaccine development. Peripheral immunogenicity is frequently measured without the paired tumor-entry, disease-response, and survival-linked sampling needed to determine why immune responses do or do not become tumor control.

Beyond external biologic support, the internal robustness of this interpretation is strengthened by the overlap-aware consolidation analysis. Readiness and drag remain separable after redundancy adjustment, the priming signal retains its statistical rank independently of readiness, and the dendritic-cell vaccine tolerogenic state is measured in manufactured product rather than donor blood, making it biologically orthogonal to the PBMC-derived host-state signals. At the same time, the redundancy analysis appropriately downgrades modules whose apparent independence was largely driven by shared gene membership and redirects attention to the unresolved downstream execution phase. In that sense, the consolidation analysis functions as an organizing filter: it preserves the biology that is independently interpretable in current public data while clarifying where additional trial architecture is still required.

The endpoint-coverage framework (Table 4) makes this gap explicit. Partitions P1 through P5 - the partitions that would link RNA expression to confirmed clinical response, whether durable benefit (P1), delayed benefit (P2), immune response without tumor control (P3), complete nonresponse (P4), or benefit without measured immune activation (P5) - are all unpopulated in current public data. Among these, P3 is especially important scientifically because it represents patients who generate measurable antigen-specific T-cell responses but fail to derive disease benefit. That is the setting in which trafficking failure, tumor microenvironment suppression, antigen loss, and durability failure become directly testable rather than merely inferred.

**Table 4.**
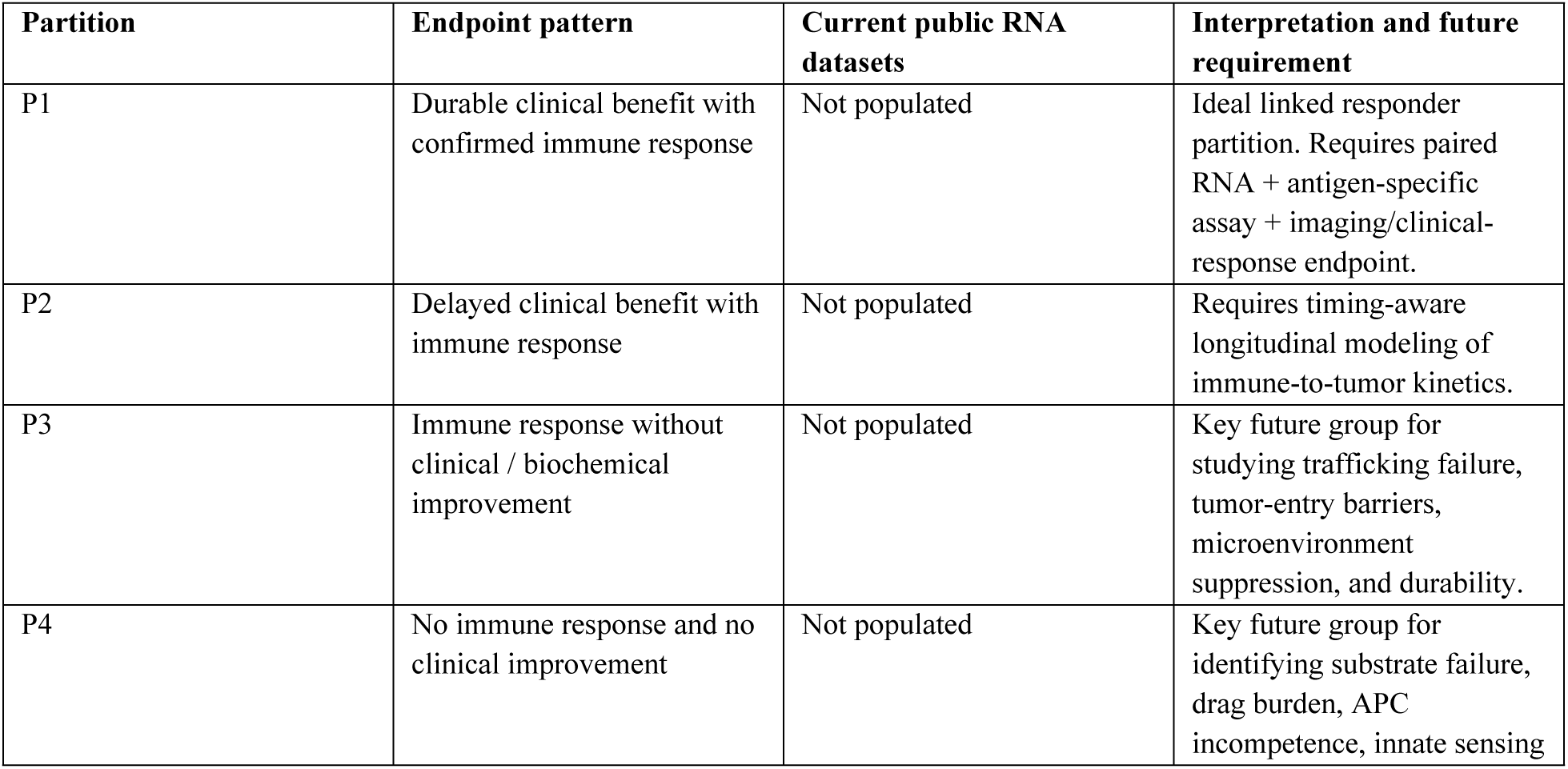
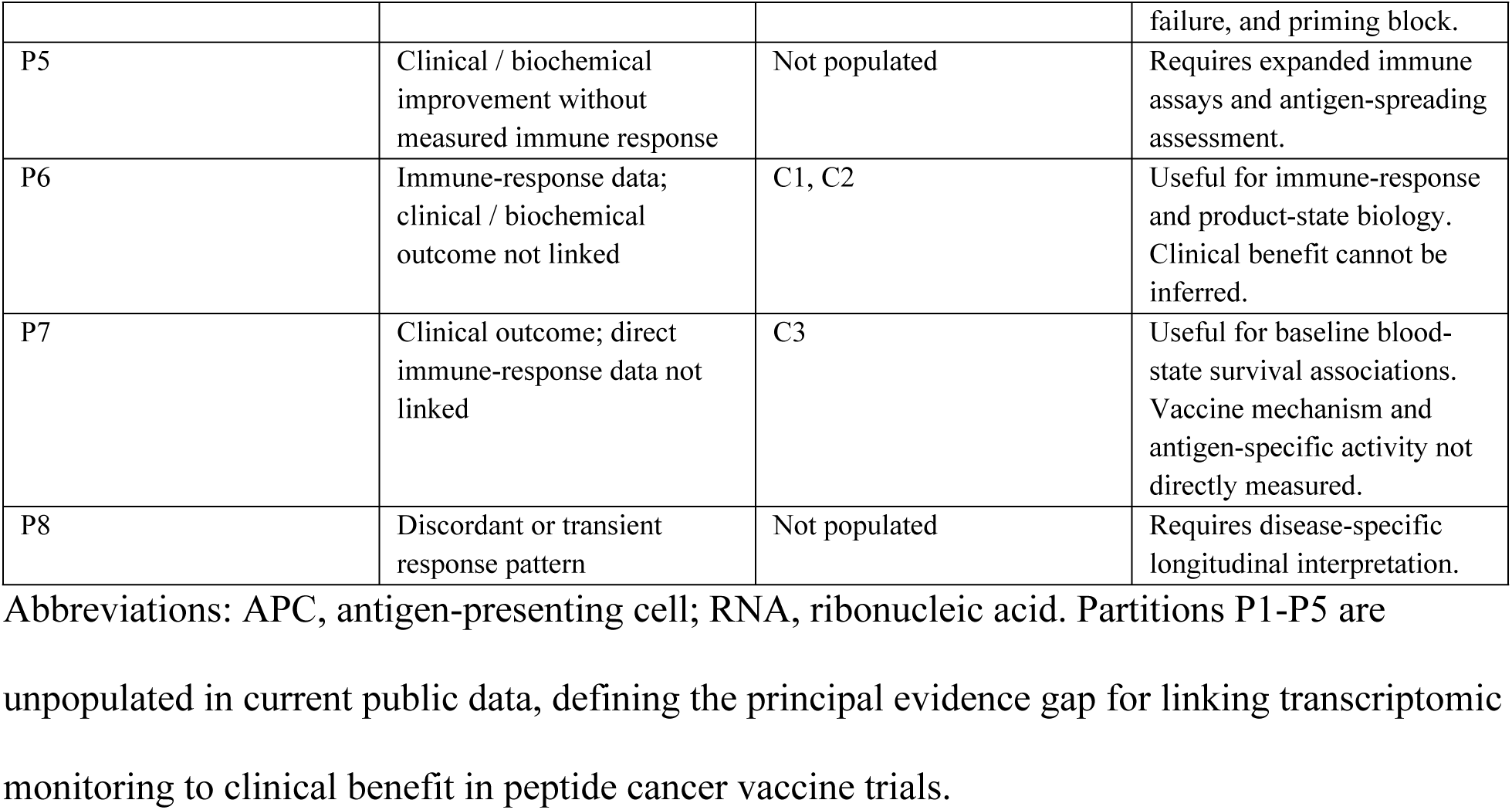
Endpoint-coverage framework for peptide vaccine transcriptomic interpretation.

A prototypical P3 patient illustrates why this partition matters. Such a patient would show ELISpot conversion and a week-2 priming signature but no prostate-specific antigen (PSA), circulating tumor DNA (ctDNA), or radiographic response. That pattern would shift the mechanistic question away from whether priming occurred and toward whether trafficking failure, tumor-entry barriers, antigen loss, intratumoral suppression, or durability failure prevented immune activity from becoming tumor control.

Closing this gap will require prospective trial design that treats transcriptomic immune monitoring, antigen-specific T-cell assays (ELISpot, T-cell receptor sequencing, multimer staining), serial imaging or PSA/cancer antigen 125 (CA125) tracking, and tumor biopsy as planned, endpoint-linked measurements rather than optional exploratory add-ons. The sequential bottleneck model provides a concrete collection framework: baseline host state (blood RNA, readiness, and drag modules), product-state quality (DC-vaccine product RNA at release), the week-2 priming window (blood RNA, priming/costimulation module), downstream trafficking and cytotoxic markers (at accessible tissue sites or through T-cell receptor repertoire sequencing), and endpoint-linked clinical data. Trials that collect these measurements within the same cohort would generate the first true P3 and P4 datasets and, with them, the first practical opportunity to dissect the immunogenicity-to-benefit gap mechanistically.

### DC product quality as a controllable pre-injection variable

The product-state finding in C2 provides a complementary example of a phase that is both biologically meaningful and operationally controllable. It connects directly to a documented manufacturing challenge: monocyte-derived dendritic-cell cultures can undergo tolerogenic drift under suboptimal maturation conditions, and this drift may not be reliably detected by standard phenotypic release criteria [7,38]. Potential contributors include incomplete Toll-like receptor-driven maturation, residual monocyte contamination, prostaglandin E2 exposure, and suboptimal cytokine conditions. CD83, CD86, and HLA-DR surface expression may appear adequate even when a product retains IL-10-producing, CD14-positive, or CD163-positive myeloid-lineage content. In C2, the direction of IL10, CD163, CD14, S100A8/S100A9, S100A12, MMP14, and TGFBI is therefore biologically coherent with known tolerogenic/myeloid drift, although the n=4 strong-responder group requires that this remain a hypothesis-generating product-state observation rather than a definitive manufacturing rule.

A transcriptional product-quality measure anchored by genes in the tolerogenic panel (CD14, CD163, IL10, MMP14, TGFBI) could serve as an orthogonal potency candidate at product release, flagging lots with high tolerogenic burden before administration. The broader principle is familiar from other cellular-therapy settings, including CAR-T cell manufacturing, where starting-cell and product-state features can influence in vivo functional performance beyond surface phenotype alone [39]. The C2 data support evaluating a similar product-state transcriptomic release metric in future dendritic-cell vaccine studies, not adopting one clinically from this small retrospective cohort.

### Candidate monitoring panels and early-futility design

Because phases 1 through 3 are measurable in current public data, they also provide the most practical starting point for structured monitoring in future trials. Taken together, these host-state, product-state, and priming findings define a practical monitoring vocabulary for future studies. Supplementary Table S6 lists candidate gene panels derived from the locked module sets as potential future monitoring tools. These are explicitly not validated biomarkers or clinical recommendations. Rather, they are structured hypotheses derived from signals that show evidence in public data and remain biologically coherent within the phase-resolved framework. Their value is to support prospective testing of each phase of the sequential model independently and to allow futility rules to be prespecified on mechanistic rather than purely empirical grounds. For clinical trial design, the sequential model implies that phase-specific transcriptomic signals could support early futility assessment: a patient with extreme erythroid/inflammatory drag at baseline and persistently low priming signal at week 2 has not cleared the first two measurable gates, and continuing vaccination alone without addressing those bottlenecks may be unlikely to produce immune benefit. Conversely, patients who clear both baseline and priming gates without showing tumor response constitute the most informative group for studying downstream execution failure and represent the priority population for add-on immunotherapy strategies targeting trafficking, tumor entry, or microenvironment suppression. This framing is intended for prospective testing and monitoring design, not for immediate clinical decision-making from the present retrospective public cohorts.

### Limitations

The principal limitation is the public data structure itself. Each cohort provides a partial, non-overlapping window into the vaccine-response cascade, and the cohorts differ in tumor type, vaccination platform, adjuvant, and patient population. This limits direct quantitative comparison across evidence partitions even as it allows cross-context evaluation of phase-aligned biology.

The C2 cohort is small (n=18 product samples, n=4 strong responders), so product-state findings are hypothesis-generating only despite exact rank-test support across the focused panel. The null week-2 paired-delta result in C1 leaves unresolved whether the cross-sectional priming signal reflects vaccine-driven induction or pre-existing biological differences that become more apparent after vaccination. The erythroid/inflammatory-drag interpretation is mechanistically coherent but relies on bulk RNA from mixed PBMC preparations and therefore requires cellular-resolution confirmation using single-cell RNA-seq, CITE-seq, or sorted-population profiling, ideally paired with hemoglobin, reticulocyte, iron/hepcidin, inflammatory-marker, treatment-history, disease-burden, and processing metadata. Finally, although all module gene sets were prespecified, the gene-set choices still reflect biological priors that require prospective validation in endpoint-linked cohorts.

## Conclusion

In summary, public peptide-vaccine transcriptomic data are consistent with a phase-resolved model with three measurable response-linked layers: baseline host immune competence (immune readiness versus erythroid/inflammatory drag), dendritic-cell product tolerogenicity, and early post-vaccine T-cell priming and costimulation. These layers are biologically grounded and directionally consistent across independent cohorts with distinct endpoints, antigens, platforms, and tumor types. The erythroid/inflammatory-drag signal, in particular, recurs across both immunogenicity and survival endpoints and therefore identifies an adverse host-state burden that warrants prospective evaluation.

The transition from peripheral immunogenicity to tumor control - the phase most critical for clinical development - cannot yet be mapped from available public data. Future prospective trials should therefore collect linked transcriptomic, immune-assay, imaging, tissue/tumor, and clinical endpoints within the same patients over time. That integrated design is the clearest next step toward understanding, predicting, and improving the clinical efficacy of peptide cancer vaccines.

### Data availability statement

The public datasets analyzed in this study are available in the Gene Expression Omnibus under accession numbers GSE278476, GSE85698, and GSE53922. Processed module-score tables, model-output tables, focused patient-level C2 expression values, prespecified gene-set definitions, supplementary methods audit files, and relative-path manuscript-level reproducibility scripts supporting this study are available from Zenodo at https://doi.org/10.5281/zenodo.20297767.

## Acknowledgments

The author gratefully acknowledges the investigators who generated, curated, and deposited the public Gene Expression Omnibus datasets used in this study.

## Supporting information

S1 Table. Supplementary Tables S1-S9. Includes prespecified module gene sets and membership, full module statistics, top baseline gene ordinal associations, erythroid enrichment/proxy checks and adjusted model refits, per-sample module scores, candidate monitoring panels, C1 covariate-adjusted permutation sensitivity, C2 exact rank-test focused-panel sensitivity, and C3 recomputed Cox checks from supporting scores.

S1 File. Supplementary Code and Audit archive. Includes relative-path manuscript-level reproducibility scripts, processed tables, prespecified gene-set definitions, focused patient-level C2 expression values, C1 sample-availability/internal-batch summaries, C2 leave-one-strong-responder sensitivity, C1/C3 joint B01+B02 sensitivity, model diagnostics, and methods audit files.

## References

1. Buonaguro L. Peptide-based vaccine for cancer therapies. Front Immunol. 2023;14:1210044. doi:10.3389/fimmu.2023.1210044. PMID:37654484. URL: https://www.frontiersin.org/journals/immunology/articles/10.3389/fimmu.2023.1210044/full

2. Fan T, et al. Therapeutic cancer vaccines: advancements, challenges, and prospects. Signal Transduct Target Ther. 2023;8(1):450. doi:10.1038/s41392-023-01674-3. PMID:38086815. URL: https://www.nature.com/articles/s41392-023-01674-3

3. Taylor-Papadimitriou J, Burchell JM, Graham R, Beatson R. Latest developments in MUC1 immunotherapy. Biochem Soc Trans. 2018;46(3):659–668. doi:10.1042/BST20170400. URL: https://portlandpress.com/biochemsoctrans/article/46/3/659/67374/Latest-developments-in-MUC1-immunotherapy

4. Gao T, Cen Q, Lei H. A review on development of MUC1-based cancer vaccine. Biomed Pharmacother. 2020;132:110888. doi:10.1016/j.biopha.2020.110888. PMID:33113416. URL: https://www.sciencedirect.com/science/article/pii/S0753332220310805

5. Lakshminarayanan V, et al. Immune recognition of tumor-associated mucin MUC1 is achieved by a fully synthetic aberrantly glycosylated MUC1 tripartite vaccine. Proc Natl Acad Sci U S A. 2012;109(1):261–266. doi:10.1073/pnas.1115166109. PMID:22171012. URL: https://pmc.ncbi.nlm.nih.gov/articles/PMC3252914/

6. Schoen RE, et al. Randomized, double-blind, placebo-controlled trial of MUC1 peptide vaccine for prevention of recurrent colorectal adenoma. Clin Cancer Res. 2023;29(9):1678–1688. doi:10.1158/1078-0432.CCR-22-3168. PMID:36892581. URL: https://pubmed.ncbi.nlm.nih.gov/36892581/

7. Castiello L, Sabatino M, Ren J, et al. Expression of CD14, IL10, and tolerogenic signature in dendritic cells inversely correlate with clinical and immunologic response to TARP vaccination in prostate cancer patients. Clin Cancer Res. 2017;23(13):3352–3364. doi:10.1158/1078-0432.CCR-16-2199. URL: https://pmc.ncbi.nlm.nih.gov/articles/PMC5496805/

8. Araki H, Pang X, Komatsu N, et al. Haptoglobin promoter polymorphism rs5472 as a prognostic biomarker for peptide vaccine efficacy in castration-resistant prostate cancer patients. Cancer Immunol Immunother. 2015;64(12):1565–1573. doi:10.1007/s00262-015-1756-7. PMID:26428930. URL: https://pubmed.ncbi.nlm.nih.gov/26428930/

9. Gene Expression Omnibus. GSE53922: baseline PBMC expression profiles from patients receiving personalized peptide vaccination for castration-resistant prostate cancer. URL: https://www.ncbi.nlm.nih.gov/geo/query/acc.cgi?acc=GSE53922

10. Cameron CM, et al. Pre-vaccination transcriptomic profiles of immune responders to the MUC1 peptide vaccine for colon cancer prevention. Front Immunol. 2024;15:1437391. doi:10.3389/fimmu.2024.1437391. URL: https://www.frontiersin.org/journals/immunology/articles/10.3389/fimmu.2024.1437391/full

11. Gene Expression Omnibus. GSE278476: MUC1 peptide vaccine PBMC RNA-seq data. URL: https://www.ncbi.nlm.nih.gov/geo/query/acc.cgi?acc=GSE278476

12. Gene Expression Omnibus. GSE85698: dendritic-cell vaccine preparation expression data from a TARP peptide vaccine trial. URL: https://www.ncbi.nlm.nih.gov/geo/query/acc.cgi?acc=GSE85698

13. Barbie DA, Tamayo P, Boehm JS, et al. Systematic RNA interference reveals that oncogenic KRAS-driven cancers require TBK1. Nature. 2009;462(7269):108–112. doi:10.1038/nature08460. PMID:19847166. URL: https://www.nature.com/articles/nature08460

14. Aran D, Hu Z, Butte AJ. xCell: digitally portraying the tissue cellular heterogeneity landscape. Genome Biol. 2017;18(1):220. doi:10.1186/s13059-017-1349-1. PMID:29141660. URL: https://genomebiology.biomedcentral.com/articles/10.1186/s13059-017-1349-1

15. Mabbott NA, Baillie JK, Brown H, Freeman TC, Hume DA. An expression atlas of human primary cells: inference of gene function from coexpression networks. BMC Genomics. 2013;14:632. doi:10.1186/1471-2164-14-632. PMID:24053356. URL: https://pubmed.ncbi.nlm.nih.gov/24053356/

16. Benjamini Y, Hochberg Y. Controlling the false discovery rate: a practical and powerful approach to multiple testing. J R Stat Soc Series B. 1995;57(1):289–300. doi:10.1111/j.2517-6161.1995.tb02031.x. URL: https://rss.onlinelibrary.wiley.com/doi/10.1111/j.2517-6161.1995.tb02031.x

17. Bradley LM, Haynes L, Swain SL. IL-7: maintaining T-cell memory and achieving homeostasis. Trends Immunol. 2005;26(3):172–176. doi:10.1016/j.it.2005.01.004. PMID:15745860. URL: https://pubmed.ncbi.nlm.nih.gov/15745860/

18. Carrette F, Surh CD. IL-7 signaling and CD127 receptor regulation in the control of T cell homeostasis. Semin Immunol. 2012;24(3):209–217. doi:10.1016/j.smim.2012.04.010. PMID:22551764. URL: https://pubmed.ncbi.nlm.nih.gov/22551764/

19. Drake A, Kaur M, Iliopoulou BP, Phennicie RT, Hanson A, et al. Interleukins 7 and 15 maintain human T cell proliferative capacity through STAT5 signaling. PLoS One. 2016;11(11):e0166280. doi:10.1371/journal.pone.0166280. URL: https://journals.plos.org/plosone/article?id=10.1371/journal.pone.0166280

20. Zhao L, He R, Long H, et al. Late-stage tumors induce anemia and immunosuppressive extramedullary erythroid progenitor cells. Nat Med. 2018;24(10):1536–1544. doi:10.1038/s41591-018-0205-5. URL: https://pmc.ncbi.nlm.nih.gov/articles/PMC6211844/

21. Grzywa TM, Justyniarska M, Nowis D, Golab J. Tumor immune evasion induced by dysregulation of erythroid progenitor cells development. Cancers (Basel). 2021;13(4):870. doi:10.3390/cancers13040870. PMID:33669537. URL: https://pubmed.ncbi.nlm.nih.gov/33669537/

22. Long H, Jia Q, Wang L, et al. Tumor-induced erythroid precursor-differentiated myeloid cells mediate immunosuppression and curtail anti-PD-1/PD-L1 treatment efficacy. Cancer Cell. 2022;40(6):674–693.e7. doi:10.1016/j.ccell.2022.04.018. PMID:35594863. URL: https://www.sciencedirect.com/science/article/pii/S1535610822002112

23. Cheng P, Corzo CA, Luetteke N, et al. Inhibition of dendritic cell differentiation and accumulation of myeloid-derived suppressor cells in cancer is regulated by S100A9 protein. J Exp Med. 2008;205(10):2235–2249. doi:10.1084/jem.20080132. PMID:18809714. URL: https://rupress.org/jem/article/205/10/2235/47046/Inhibition-of-dendritic-cell-differentiation-and

24. Sinha P, Okoro C, Foell D, Freeze HH, Ostrand-Rosenberg S, Srikrishna G. Proinflammatory S100 proteins regulate the accumulation of myeloid-derived suppressor cells. J Immunol. 2008;181(7):4666–4675. doi:10.4049/jimmunol.181.7.4666. PMID:18802069. URL: https://pubmed.ncbi.nlm.nih.gov/18802069/

25. Ali AK, Nandagopal N, Lee SH. IL-15-PI3K-AKT-mTOR: a critical pathway in the life journey of natural killer cells. Front Immunol. 2015;6:355. doi:10.3389/fimmu.2015.00355. PMID:26257729. URL: https://pubmed.ncbi.nlm.nih.gov/26257729/

26. Nandagopal N, Ali AK, Komal AK, Lee SH. The critical role of IL-15-PI3K-mTOR pathway in natural killer cell effector functions. Front Immunol. 2014;5:187. doi:10.3389/fimmu.2014.00187. URL: https://www.frontiersin.org/journals/immunology/articles/10.3389/fimmu.2014.00187/full

27. Wang X, et al. Transcription factors associated with IL-15 cytokine signaling during NK cell development. Front Immunol. 2021;12:610789. doi:10.3389/fimmu.2021.610789. PMID:33815365. URL: https://pubmed.ncbi.nlm.nih.gov/33815365/

28. Zhao Y, et al. IL-7: a promising adjuvant ensuring effective T cell responses and memory in combination with cancer vaccines? Front Immunol. 2022;13:1022808. doi:10.3389/fimmu.2022.1022808. PMID:36389666. URL: https://www.frontiersin.org/journals/immunology/articles/10.3389/fimmu.2022.1022808/full

29. Pellegrini M, Calzascia T, Elford AR, et al. Adjuvant IL-7 antagonizes multiple cellular and molecular inhibitory networks to enhance immunotherapies. Nat Med. 2009;15(5):528–536. doi:10.1038/nm.1953. PMID:19396174. URL: https://www.nature.com/articles/nm.1953

30. Colombetti S, Levy F, Chapatte L. IL-7 adjuvant treatment enhances long-term tumor-antigen-specific CD8+ T-cell responses after immunization with recombinant lentivector. Blood. 2009;113(26):6629–6637. doi:10.1182/blood-2008-05-155309. PMID:19383968. URL: https://ashpublications.org/blood/article/113/26/6629/26219/IL-7-adjuvant-treatment-enhances-long-term-tumor

31. Moller P, Sun Y, Dorbic T, et al. Vaccination with IL-7 gene-modified autologous melanoma cells can enhance the anti-melanoma lytic activity in peripheral blood of patients with a good clinical performance status: a clinical phase I study. Br J Cancer. 1998;77(11):1907–1916. doi:10.1038/bjc.1998.317. PMID:9667667. URL: https://pmc.ncbi.nlm.nih.gov/articles/PMC2150323/

32. Pachynski RK, Morishima C, Szmulewitz R, et al. IL-7 expands lymphocyte populations and enhances immune responses to sipuleucel-T in patients with metastatic castration-resistant prostate cancer (mCRPC). J Immunother Cancer. 2021;9(8):e002903. doi:10.1136/jitc-2021-002903. PMID:34452927. URL: https://jitc.bmj.com/content/9/8/e002903.long

33. Lui G, Minnar CM, Soon-Shiong P, Schlom J, Gameiro SR. Exploiting an interleukin-15 heterodimeric agonist (N803) for effective immunotherapy of solid malignancies. Cells. 2023;12(12):1611. doi:10.3390/cells12121611. PMID:37371081. URL: https://pmc.ncbi.nlm.nih.gov/articles/PMC10297013/

34. ClinicalTrials.gov. NCT05419011: Testing a combination of vaccines for cancer prevention in Lynch syndrome; Adenovirus 5 CEA/MUC1/Brachyury Vaccine Tri-Ad5 and Nogapendekin Alfa (N-803). URL: https://clinicaltrials.gov/study/NCT05419011

35. National Cancer Institute. Phase I/II study of SX-682, TriAdeno CEA/MUC1/Brachyury vaccine, retifanlimab, and IL-15 agonist N-803 (STAR15) for metastatic colorectal cancer; NCT06149481. URL: https://www.cancer.gov/research/participate/clinical-trials/intervention/adenovirus-5-cea-muc1-brachyury-vaccine-tri-ad5

36. Fraenkel PG. Anemia of inflammation. Med Clin North Am. 2017;101(2):285–296. doi:10.1016/j.mcna.2016.09.005. URL: https://pmc.ncbi.nlm.nih.gov/articles/PMC5308549/

37. Marques O, Weiss G, Muckenthaler MU. The role of iron in chronic inflammatory diseases: from mechanisms to treatment options in anemia of inflammation. Blood. 2022;140(19):2011–2023. doi:10.1182/blood.2021013472. URL: https://www.sciencedirect.com/science/article/pii/S0006497122010862

38. Butterfield LH. Dendritic cell-based vaccines: barriers and opportunities. Future Oncol. 2012;8(10):1273–1299. doi:10.2217/fon.12.125. PMID:23130928. URL: https://pmc.ncbi.nlm.nih.gov/articles/PMC4260651/

39. Fraietta JA, Lacey SF, Orlando EJ, et al. Determinants of response and resistance to CD19 chimeric antigen receptor (CAR) T cell therapy of chronic lymphocytic leukemia. Nat Med. 2018;24(5):563–571. doi:10.1038/s41591-018-0010-1. PMID:29713085. URL: https://pmc.ncbi.nlm.nih.gov/articles/PMC6117613/

